# High-precision neural stimulation by a highly efficient candle soot fiber optoacoustic emitter

**DOI:** 10.1101/2022.07.30.502146

**Authors:** Guo Chen, Linli Shi, Lu Lan, Runyu Wang, Yueming Li, Zhiyi Du, Mackenzie Hyman, Ji-Xin Cheng, Chen Yang

## Abstract

Highly precise neuromodulation with a high efficacy poses great importance in neuroscience. Here, we developed a candle soot fiber optoacoustic emitter (CSFOE), capable of generating a high pressure of over 10 MPa, enabling highly efficient neuromodulation in vitro. The design of the fiber optoacoustic emitter, including the choice of the material and the thickness of the layered structure, was optimized in both simulations and experiments. The optoacoustic conversion efficiency of the optimized CSFOE was found to be ten times higher than the other carbon-based fiber optoacoustic emitters. Driven by a single laser, the CSFOE can perform dual-site optoacoustic activation of neurons, confirmed by calcium (Ca) imaging. Our work opens potential avenues for more complex and programmed control in neural circuits using a simple design for multisite neuromodulation in vivo.

## 1 Introduction

Highly precise neural modulation is of great importance in neuroscience, as the firing of specific neuronal populations in the brain could alter the behavior of animals which could serve as a novel tool for studying neural pathways in disease and health. Sophisticated control of neuronal circuits and brain functions requires stimulating multiple functional regions at high spatial resolution. For example, a previous study by Li, et al used two ultrasound transducers to stimulate primary somatosensory cortex barrel field (S1BF) of a free moving mouse and successfully controlled the head turning direction of the mouse by applying stimuli at different position (Li et al., 2019). Among the current neuromodulation platforms, electrical neuron stimulation has been proven to be efficient and allows for deep brain stimulation, while it provides a limited spatial resolution of millimeters in vivo, due to electric current spread (Boon et al., 2007). Optogenetics neural stimulation with single neuron resolution has been shown as a powerful tool in fundamental studies, but the requirement of viral infection makes it challenging to apply to human brains (Boyden et al., 2005). Transcranial magnetic stimulation (TMS) and transcranial direct current stimulation (tDCS) are capable of non-invasive transcranial neuromodulation, while suffering from the resolution at the centimeter level(Rosa and Lisanby, 2012;Davidson et al., 2020). Infrared neuron stimulation (INS) takes advantage of the near-infrared absorption of water to generate heat for neuron stimulation. However, the thermal toxicity and potential tissue damage is a concern in real clinical scenarios (Cayce et al., 2014;Zhu et al., 2022). Focused ultrasound is an emerging non-invasive modality with deep penetration depth in tissue (Beisteiner et al., 2020;Bobola et al., 2020;Brinker et al., 2020). It has a spatial resolution limited by the acoustic wave’s diffraction, therefore it is challenging for low-frequency (< 1 MHz) ultrasound to reach submillimeter level. New technologies and methods are still being sought for precise and non-genetic neural stimulation.

Recently, our team developed a miniaturized fiber-optoacoustic converter (FOC) converting pulsed laser into ultrasound (Jiang et al., 2020). FOC succeeded in spatially confined neural stimulation of mouse brain and modulation of motor activity in vivo. It was found that, for successful FOC based optoacoustic neural stimulation, a pressure of around 0.5 MPa is needed (Jiang et al., 2020). Typical FOC generate a pressure of 0.48 MPa upon laser pulse energy of 14.5 μJ, with an estimated photoacoustic conversion efficiency of 1374 Pa m^2^/J. Considering the typical energy and repetition rate of nanosecond lasers, the low conversion efficiency of FOC limits its application in multi-site stimulation. Thus, new fiber optoacoustic emitters with a higher conversion efficiency are needed to enable multisite optoacoustic neuromodulation.

According to a simplified model of optoacoustic generation, the output optoacoustic pressure is related to the laser fluence (*F*), the absorption coefficient (*α*) and the thermal expansion coefficient (*β*) (Xu and Wang, 2006). The pressure generated can be calculated following the equation below:

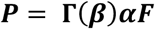

where the Grüneisen parameter (*┌*) is a function of the thermal expansion coefficient *β*. To improve the optoacoustic conversion efficiency, materials with greater light absorption and larger thermal expansion coefficients will be preferable choices. Previously, many materials have been studied for efficient optoacoustic conversion, including metal, carbon material, etc. Metals, in the form of gold nanoparticles (Wu et al., 2011;Wu et al., 2012;Tian et al., 2013;Wu et al., 2013;Zou et al., 2014) and Cr and Ti films(Lee and Guo, 2017) were used due to their high absorption coefficient. However, the high light reflection by metal films and scattering of metal nanoparticles limits the energy conversion efficiency. Different carbon materials have also been studied, including carbon nanoparticle (CNP) (Biagi et al., 2001), carbon nanotube (CNT) (Won Baac et al., 2010;Colchester et al., 2014;Baac et al., 2015;Alles et al., 2016;Noimark et al., 2016;Moon et al., 2017;Poduval et al., 2017;Shi et al., 2020;Thompson et al., 2022), graphite (Jiang et al., 2020) and candle soot films(CS) (Chang et al., 2015;Huang et al., 2016;Chang et al., 2018). Among these, candle soot stands out for its high light absorption coefficient, low interfacial thermal resistance, and easy fabrication process. Direct comparison of optoacoustic conversion efficiency among CS, CNT and CNP showed CS can generate a pressure six times higher than that generated by the other two materials (Chang et al., 2018). The CS layer deposited onto polydimethylsiloxane (PDMS), a material with high thermal expansion coefficient(Wolf et al., 2018), forms a diffused mixture – an excellent choice for highly efficient optoacoustic generation.

In this work, we developed a candle soot-based fiber optoacoustic emitter (CSFOE) for the first time. COMSOL simulation was used to simulate the optoacoustic generation process of a CSFOE. We optimized the design of the CSFOE by identifying the optimal thickness of the CS layer through simulation. A CSFOE with a CS layer of an optimal ∼ 10 μm thickness was found to achieve the highest peak-to-peak pressure. Experimentally, we fabricated CSFOEs with controlled thickness of candle soot layers in the range of 1 μm to 60 μm. By comparing their optoacoustic performance, we confirmed that the optimal thickness of the CS layer is 10 μm, consistent with the simulation prediction. The maximum optoacoustic pressure reached ∼10 MPa, which is 9.6 times larger compared with that generated by FOC (Jiang et al., 2020;Shi et al., 2021). The application of CSFOE to non-genetic optoacoustic neural stimulation was demonstrated in GCaMP labeled neuron culture. Successful high precision activation of neurons confined in an area of 200 μm was verified by calcium imaging. Significantly, we demonstrated dual-site optoacoustic neuron stimulation driven by a single laser utilizing the high optoacoustic conversion efficiency of CSFOE. The highly localized ultrasound field generated by each CSFOE allows the two stimulated sites to be sub-millimeter apart. Our work opens up potentials for complex and programmed control in neural circuits using a simple design for multisite neuromodulation.

## 2 Material and Methods

### 2.1 Simulation of the ultrasound field generated from CSFOE

The ultrasound field generated by CSFOE was simulated by COMSOL Multiphysics 5.4. The CSFOE was modeled by a 2D axisymmetric model with a double-layer structure, including an absorber layer of CS and PDMS mixture and a pure PDMS layer. The backing material was set to be a fiber (SiO_2_). Radiative Beam in Absorbing Media module was used to simulate the absorption of the CSNP-PDMS mixture layer when being applied a nanosecond pulsed laser. The Heat Transfer and Solid Mechanics modules were used to model the thermal expansion caused by the photothermal effect. The Transient Pressure Acoustic module converted the thermal expansion of the absorber into the acoustic signal and simulated the propagation of ultrasound in the water medium. The absorption coefficient of the absorber layer used in the simulation was a measured value by our experiment, and all the other material parameters were set according to COMSOL’s material library database.

### 2.2 Fabrication of CSFOE

A schematic of the CSFOE is shown in **Figure 1a**. A flame from a paraffin wax candle served as the source of the candle soot. To fabricate the CSFOE, the tip of a polished multimode optical fiber (with a 200 μm diameter (200EMT, Thorlabs)) was placed into the center of the flame for three to five seconds. This step was repeated until the optical fiber was fully coated with flame synthesized candle soot. To prepare the PDMS, the silicone elastomer (Sylgard 184, Dow Corning Corporation, USA) was carefully dispensed into a container to minimize air entrapment. Then, the curing agent was added for a weight ratio of ten to one (silicone elastomer to curing agent). A nanoinjector deposited the PDMS onto the tip of the candle-soot coated fiber. The position of the fiber and the nanoinjector were both controlled by 3D manipulators for precise alignment, and the PDMS coating process was monitored under a lab-made microscope in real time. The coated fiber was stored overnight in a temperature-controlled environment, to allow the PDMS to cure and fully diffuse into the porous structure of the candle soot.

**Figure 1.**
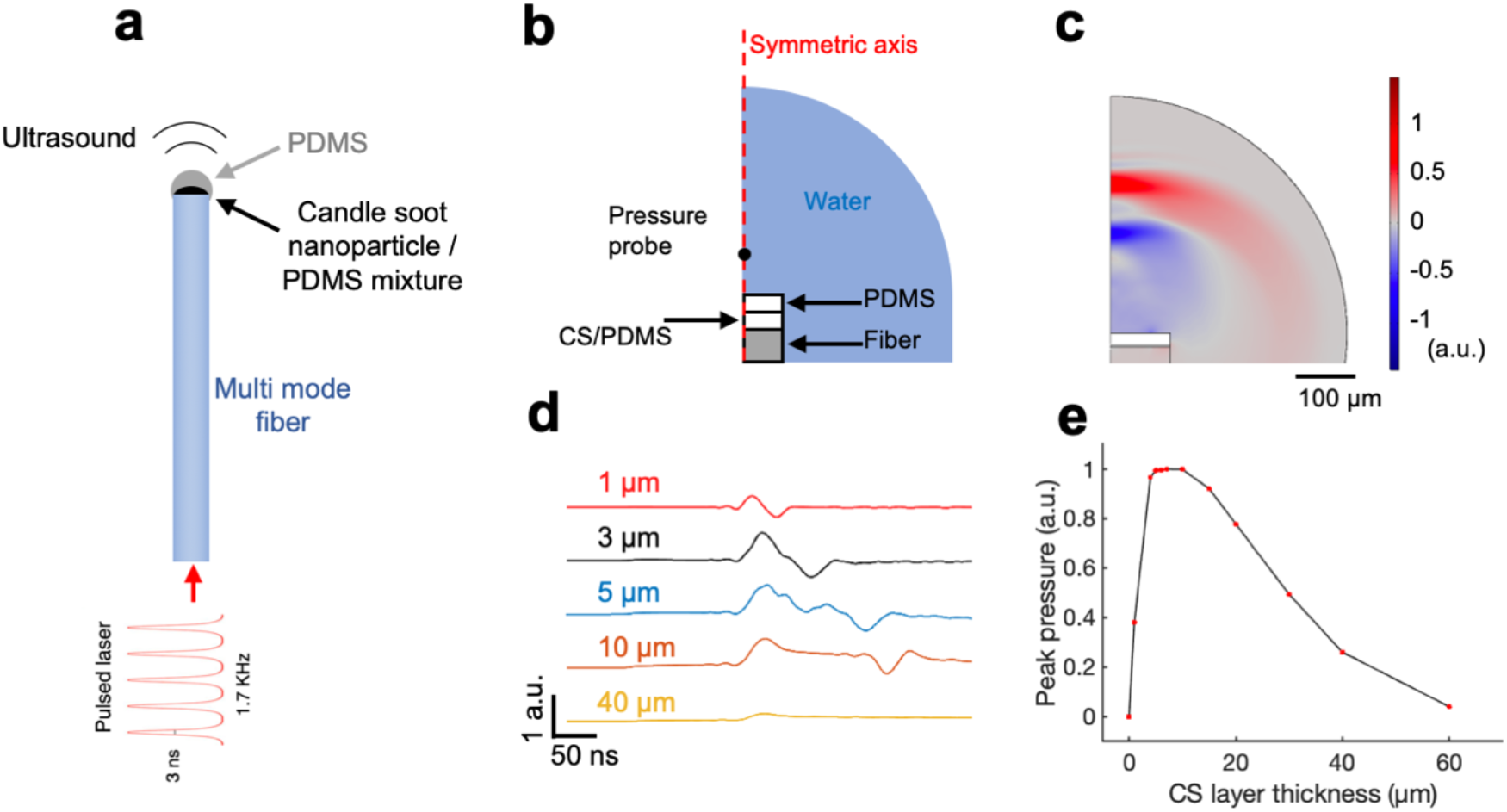
COMSOL simulation of CSFOE performance. (a) Schematic of CSFOE. (b) Illustration of the CSFOE model used in simulation. Not to scale. (c) Representative ultrasound waveform, simulated at t=400 ns under an input of a 3 ns pulsed laser. (d) Acoustic waveforms simulated at different thicknesses of the CS layer. (e) Peak-to-peak acoustic pressure plotted as a function of candle soot layer thickness.

### 2.3 Characterization of absorption ability of CSNP

The absorption of CS with different deposition thickness was measured with a photodiode (Thorlabs, USA). Different thicknesses of CS were controlled by the deposition durations. The fiber was connected to a Q-switched 1030-nm nanosecond laser (Bright Solution, Inc. Calgary Alberta, CA), and the transmission power was detected by a laser diode.

### 2.4 Characterization of optoacoustic signals

The amplitude of the CSFOE-generated acoustic wave was measured using a needle hydrophone with a 40 μm core sensor (Precision Acoustics, UK). A digital oscilloscope (DSO6014A, Agilent Technologies, CA, USA) recorded the electrical signal from the hydrophone. A four-axis micro-manipulator (MC1000e controller with MX7600R motorized manipulator, Siskiyou Corporation, OR, USA), with a resolution of 0.2 μm, controlled the distance between the CSFOE tip and the hydrophone, which was incremented from 0 to 400 μm. The distance was measured using a widefield microscope with a 10× objective. The CSFOE tip and the hydrophone tip were both immersed in a drop of degassed water placed on a glass microscope slide. The CSFOE was connected to a Q-switched 1030-nm nanosecond laser (Bright Solution, Inc. Calgary Alberta, CA) with a laser pulse energy of 56 μJ. The setup of the measurement is shown in **Figure 2d**. The acoustic pressure values were calculated based on the calibration curve obtained from the hydrophone manufacturer. The frequency data were obtained through the Fast Fourier Transform (FFT) using MATLAB R2020a. For visualizing the acoustic wavefront, a Q-switch Nd: YAG laser (Quantel Laser CFR ICE450) was used to deliver 8 ns pulses to the CSFOE. The generated acoustic signal was capture by a 1×128 linear transducer array (L22-14, Verasonics Inc.) and processed by an ultrasound imaging system (Vantage128, Verasonics).

**Figure 2.**
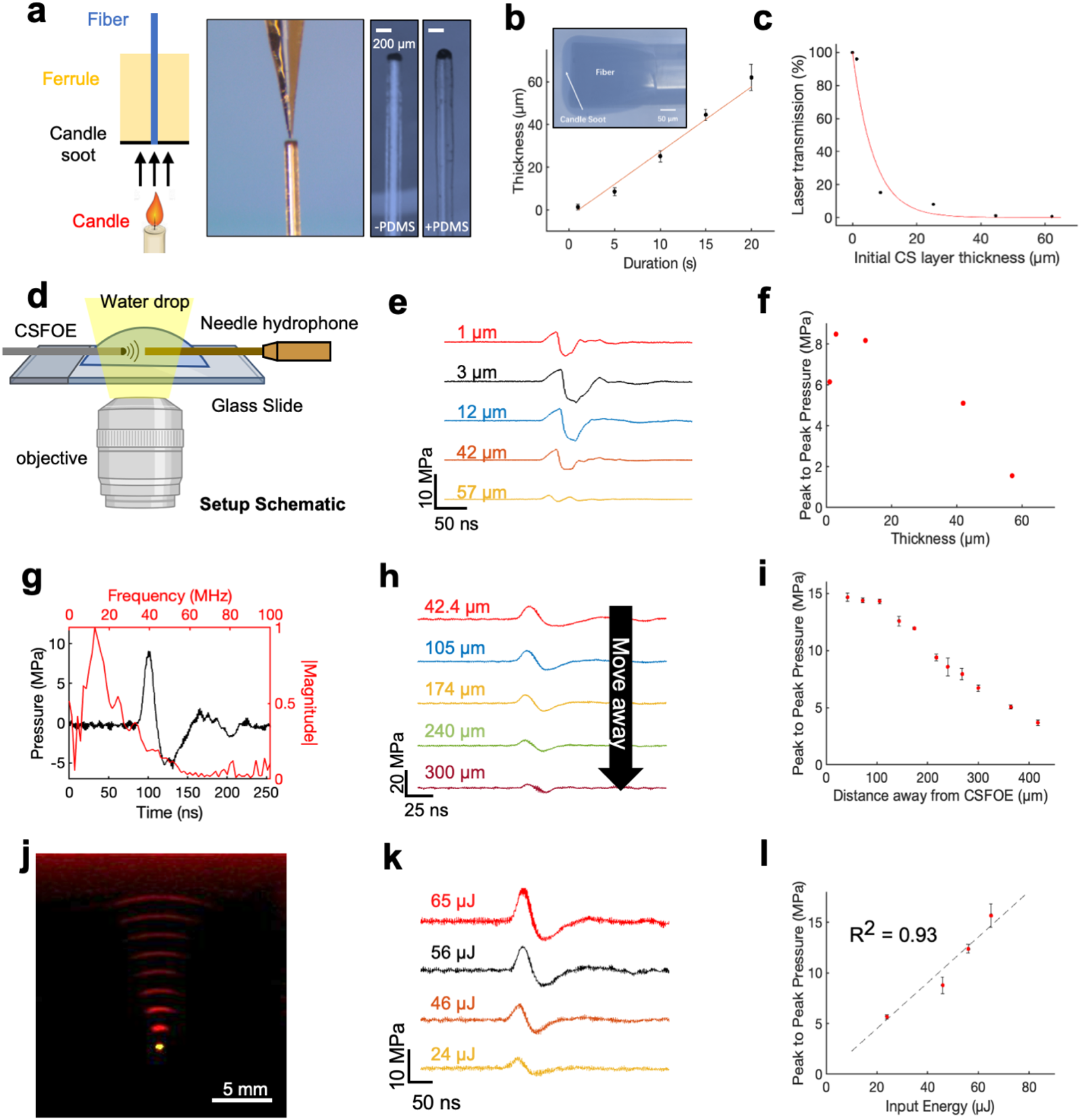
Fabrication and characterization of CSFOE. (a) Key steps of CSFOE fabrication. (left) Candle soot deposition on an optical fiber tip. (middle) PDMS coating on the surface of the CS layer using a nanoinjector. (Right) Images of samples after CS deposition and PDMS coating, respectively. Scale bars: 200 μm. (b) Thickness of CS layer obtained as a function of deposition duration. Inset: representative image of a fiber coated with candle soot. (c) Transmission ratio plotted as a function of the thickness of CS layer. (d) Schematic of experimental configuration of photoacoustic signal measurement using a 40 μm needle hydrophone. (e) Acoustic signal of CSFOE as a function of the candle soot layer thickness detected by the hydrophone. Laser condition: 1030nm, 1.7 kHz repetition rate, 56 μJ per pulse. (f) Peak to peak pressure plotted as a function of the thickness of CS under the same laser condition as (e). (g) Representative photoacoustic waveform (black) detected by the hydrophone and its FFT frequency spectrum (red). (h, i) Acoustic signal and peak-to-peak pressure generated by CSFOE detected at different distances from the CSFOE tip. Each data point was an average of three trials. (j) Photoacoustic signal propagation in the medium detected by a linear transducer array. Fiber tip (yellow), PA waveform (red). (k, l) Photoacoustic waveforms and peak-to-peak pressures measured at different laser pulse input. Each data point was an average of three trials.

### 2.5 Embryonic neuron culture

Primary cortical neuron cultures were derived from Sprague-Dawley rats. Cortices were dissected from embryonic day 18 (E18) rats of either sex and then digested in papain (0.5 mg/mL in Earle’s balanced salt solution) (Thermo Fisher Scientific Inc., MA). Dissociated cells were washed with and triturated in 10% heat-inactivated fetal bovine serum (FBS, Atlanta Biologicals, GA, USA), 5% heat-inactivated horse serum (HS, Atlanta Biologicals, GA, USA), 2mM Glutamine-Dulbecco’s Modified Eagle Medium (DMEM, Thermo Fisher Scientific Inc., MA, USA), and cultured in cell culture dishes (100 mm diameter) for 30 minutes at 37 °C to eliminate glial cells and fibroblasts. The supernatant containing neurons was collected and seeded on poly-D-lysine coated cover glass and incubated in a humidified atmosphere containing 5% CO_2_ at 37 °C with 10% FBS + 5% HS + 2 mM glutamine DMEM. After 16 hours, the medium was replaced with Neurobasal medium (Thermo Fisher Scientific Inc., MA, USA) containing 2% B27 (Thermo Fisher Scientific Inc., MA, USA), 1% N2 (Thermo Fisher Scientific Inc., MA, USA), and 2mM glutamine (Thermo Fisher Scientific Inc., MA, USA). On day five, cultures were treated with 5 μM FDU (5-fluoro-2’-deox-yuridine, Sigma-Aldrich, MO, USA) to further reduce the number of glial cells. Additionally on day five, the AAV9.Syn.Flex.GCaMP6f.WPRE.SV40 virus (Addgene, MA, USA) was added to the cultures at a final concentration of 1 μl/mL for GCaMP6f expression. Half of the medium was replaced with fresh culture medium every three to four days. Cells cultured in vitro for 10–13 days were used for CSFOE stimulation experiment.

### 2.6 In vitro neurostimulation

In vitro neurostimulation experiments were performed using a Q-switched 1030-nm nanosecond laser (Bright Solution, Inc. Calgary Alberta, CA). A 3-D micromanipulator (Thorlabs, Inc., NJ, USA) was used to position the CSFOE in the cell culture dish. Calcium fluorescence imaging was performed on a lab-built wide-field fluorescence microscope based on an Olympus IX71 microscope frame with a 20× air objective (UPLSAPO20X, 0.75NA, Olympus, MA, USA), illuminated by a 470 nm LED (M470L2, Thorlabs, Inc., NJ, USA) and a dichroic mirror (DMLP505R, Thorlabs, Inc., NJ, USA). Image sequences were acquired with a scientific CMOS camera (Zyla 5.5, Andor) at 20 frames per second. Neurons expressing GCaMP6f at DIV (day in vitro) 10–13 were used for the stimulation experiment. For the tetrodotoxin (TTX) control group, tetrodotoxin citrate (ab120055, Abcam, MA, USA) was added to the culture to reach 3 μM final concentration 10 min before Calcium imaging. The fluorescence intensities, data analysis, and exponential curve fitting were analyzed using ImageJ (Fiji) and MATLAB R2020a.

### 2.7 Data analysis

Calcium images were analyzed using ImageJ. The fluorescence intensity was measured by selecting the soma. Calcium traces, acoustic waveform and temperature traces were analyzed using MATLAB R2020a. All statistical analysis was done using MATLAB R2020a. Data shown are mean ± standard deviation.

## 3 Results

### 3.1 Simulation of acoustic waveforms generated from CSFOE

To identify the optimal condition towards maximized optoacoustic conversion efficiency, we used COMSOL Multiphysics to simulate the generation and propagation of the optoacoustic signals. Taking advantage of COMSOL multiple physics field simulations, we simulated the different steps of optoacoustic generation: laser absorption, thermal expansion, and acoustic wave propagation. Since the candle soot has a very porous structure, the PDMS diffuses into the candle soot layer, forming a uniformly mixed candle soot/PDMS mixture layer(Chang et al., 2015). Therefore, we included a candle soot/PDMS mixture layer and a pure PDMS layer in the 2D axisymmetric CSFOE model built in COMSOL Multiphysics 5.4 for simulation (**Figure 1b**). A single 3 ns laser pulse was delivered through a multimode fiber (with a 200 μm core diameter) to the double layer coating on the fiber tip. **Figure 1c** shows a representative wave front of the generated ultrasound 400 ns after the onset of the laser, indicating a bipolar pressure signal generated by CSFOE (**Figure 1c**, red: positive pressure; blue: negative pressure). **Figure 1d** plots the time domain waveforms when the thickness of the CS/PDMS layer varied from 1 μm to 40 μm. In **Figure 1e**, the normalized peak-to-peak amplitude of the generated PA signal is plotted as a function of the thickness of the CS/PDMS mixture layer. The optimal thickness of the CS/PDMS layer, which generates the largest amplitude PA signal, was found to be ∼10 μm. This result is consistent with the previous work, where an optimal thickness generating the maximum pressure was also found (Chang et al., 2018).

### 3.2 Fabrication and characterization of CSFOE

To fabricate the optimal CSFOE, as guided by the simulations, we developed a two-step fabrication procedure to precisely control the CS layer thickness (**Figure 2a**). A polished multimode fiber with a 200 μm core diameter (Thorlabs) was inserted into a fiber ferrule. The fiber tip was positioned so that it was flush with the distal end of the ferrule. Then, the distal tips of both the ferrule and fiber were placed into the flame core of a paraffin wax candle, where they were fully coated with flame synthesized candle soot (**Figure 2a**, left). The key parameter to control the thickness of the CS was the coating time, which ranged between 1 to 20 s. Then, a nanoinjector was used to deposit a controlled amount of PDMS (∼0.01 μm^3^) onto the tip of the fiber coated with candle soot (**Figure 2a**, middle). The transmission images of the CS-coated fiber before and after PDMS coating are also shown in **Figure 2a** (right). When varying the CS coating time, the thickness of the CS coating was measured from the transmission images of samples before PDMS coating. **Figure 2b** plots the thickness of the CS layer measured as a function of deposition time. The CS layer thickness was linearly proportional to the deposition time, with an estimated deposition rate of 3.04 μm/s, similar to previous reports(Chang et al., 2018). Such a linear relation enables us to precisely control the thickness of the CS layer to study the PA conversion as a function of the layer thickness. Transmission of CS layers with different thicknesses were also measured (**Figure 2c**). The normalized transmission of the coating exponentially decreased as a function of the thickness. An absorption depth, the thickness when the transmission decreased to 1/e of initial transmission at the zero thickness was obtained as 6.6 μm. This measured ultrathin absorption depth indicated strong absorption of CS in NIR, enabling efficient ultrasound generation.

The characterization of the CSFOE with various CS/PDMS layer thicknesses was performed with a 40 μm needle hydrophone. A 1030 nm nanosecond pulsed laser, with a 46 μJ pulse energy was delivered to the CSFOE to generate optoacoustic signals. The acoustic signals were measured for CSFOEs where the thickness of the CS-PDMS mixture layer ranged from 1 μm to 57 μm (**Figure. 2d**). The peak-to-peak pressure is plotted as a function of the CS layer thickness in **Figure 2f**. An optimal thickness of ∼10 μm was found to generate the highest peak-to-peak pressure of 9 MPa. Notably, the experimentally measured optimal thickness and the trend between the thickness and the peak-to-peak pressure are consistent with the simulation results. Importantly, the 10 μm optimal thickness was also found to be close to the 6.6 μm absorption depth of CS-PDMS layer obtained from the absorption shown in **Figure 2c**. The greatest optoacoustic conversion efficiency occurred when the absorption layer thickness equaled the material absorption depth. In the thickness range < 10 μm, when increasing the absorption layer thickness first, the thickness at the absorption depth allowed complete optical absorption. Further increasing the thickness beyond the absorption depth (>10 μm) led to acoustic attenuation, as demonstrated in previous works(Chang et al., 2018).

Frequency characterization of the generated optoacoustic signal is shown in **Figure 2g**. The frequency spectrum of the measured acoustic waveforms after Fast Fourier Transform (FFT) exhibited a peak acoustic frequency of 12.8 MHz. This frequency was similar to previous studies on candle-soot-based optoacoustic films(Chang et al., 2018), in which a central frequency of ∼10 MHz was detected for ∼2 μm CS coating thickness. To map the propagation of the optoacoustic wave generated by the CSFOE, the pressure was measured at different distances away from the CSFOE using a 40 μm needle hydrophone as shown in **Figure 2h**. The peak-to-peak pressure of the generated ultrasound is plotted as a function of distance in **Figure 2i**. The measurements were repeated for three times and the average values were plotted. The confinement of the generated acoustic field, defined by the distance where the pressure decreases to 1/e of the initial pressure at 0 μm, was found to be ∼300 μm, approximately equal to the size of the fiber core. Such decay of optoacoustic pressure over the distance away from the CSFOE tip enables a sub-millimeter localized neuron stimulation. In addition, **Figure 2i** shows that the dependence on distance is different from the previous 1/r^2^ relation obtained in FOC. The difference is due to the fact that the ultrasound field emitted by CSFOE is at a higher frequency, therefore propagates more directionally, compared with more omnidirectional propagation of the lower frequency FOCs(Jiang et al., 2020).

The propagation of generated ultrasound can be directly visualized using an optoacoustic tomography system (**Figure 2j**). The acoustic signal was detected by a 1 × 128 linear transducer array (L22-14, Verasonics Inc.) and processed by an ultrasound imaging system (Vantage128, Verasonics). The emitted ultrasound waveform (red) obtained with a time interval of 0.5 μs and the image of the tip of the CSFOE (yellow) are overlaid in **Figure 2j**. Through the photoacoustic waveform shown in **Figure 2j**, the emission angle of CSFOE was measured to be 25.3 degrees. For FOC reported previously (Jiang et al., 2020), the emission angle was measured to be 55.1 degrees which is around twice as large. This observation also supports the more directional propagation for the CSFOE generated ultrasound field(Jiang et al., 2020).

Different laser energy inputs also resulted in varied output pressures. Using different fiber attenuators to control the laser energy input, the waveform of generated acoustic signal was measured by the needle hydrophone (**Figure. 2k**), and the peak-to-peak pressure is plotted as a function of input energy in **Figure 2l**, showing a fitting curve of P = 0.226 * E (R^2^ = 0.93, fitting coefficient of determination) and confirming the linear dependence of the pressure on the input laser energy.

Through controlling the distance away from the CFOE tip and laser energy, we can have a complete control of the generated pressure in a large range under 15 MPa for various applications. By rationale fabrication of the layered structure of CSFOE and control of PA pressure generated, CSFOE can serve as a robust device for repeatable neuromodulation and allows us to study neuron responses under different conditions.

### 3.3 CSFOE Stimulation of neurons in vitro

To confirm the stimulation function of the CSFOE, GCaMP6f-labeled primary neurons (DIV 12-14) were cultured on a glass bottom dish, and calcium imaging was performed to monitor neuronal activities. A 3 ns pulsed laser at 1030 nm with a repetition rate of 1.7 kHz was delivered to the CSFOE. The laser pulse train, with a duration of 3 ms (corresponding to 5 pulses) and pulse energy of 65 μJ, was used for CSFOE optoacoustic in vitro neural stimulation. The CSFOE was precisely controlled by a 4D micromanipulator to approach the target neurons. The distance between the neurons and the CSFOE tip was monitored to make sure neurons were within the sub-millimeter confinement area.

Representative fluorescence images of the neuron before and after stimulation are shown in **Figure 3a** and **b**. Maximum change of the fluorescence intensity is highlighted in **Figure 3c**. The dashed circles indicate the location of the CSFOE. Increase in fluorescence intensity reaching ΔF/F_0_>10% upon stimulation confirms the successful activation. This map of fluorescence changes in **Figure 3c** also in indicates that neurons within the stimulation area were successfully activated. The activation outside the stimulation area is due to networking effect (more details discussed later). To further investigate whether the CSFOE can activate neurons reliably and repeatedly, we stimulated the same area of neuron three times in four minutes (**Figure 3d)**. Repeatable stimulations were successfully observed after the laser onset at t = 5 s, 90 s and 180 s, and all show Δ*F*/*F*_0_>10%. This result clearly shows that there is no damage caused by CSFOE after stimulation and demonstrates the repeatability and safety of CSFOE stimulation.

**Figure 3.**
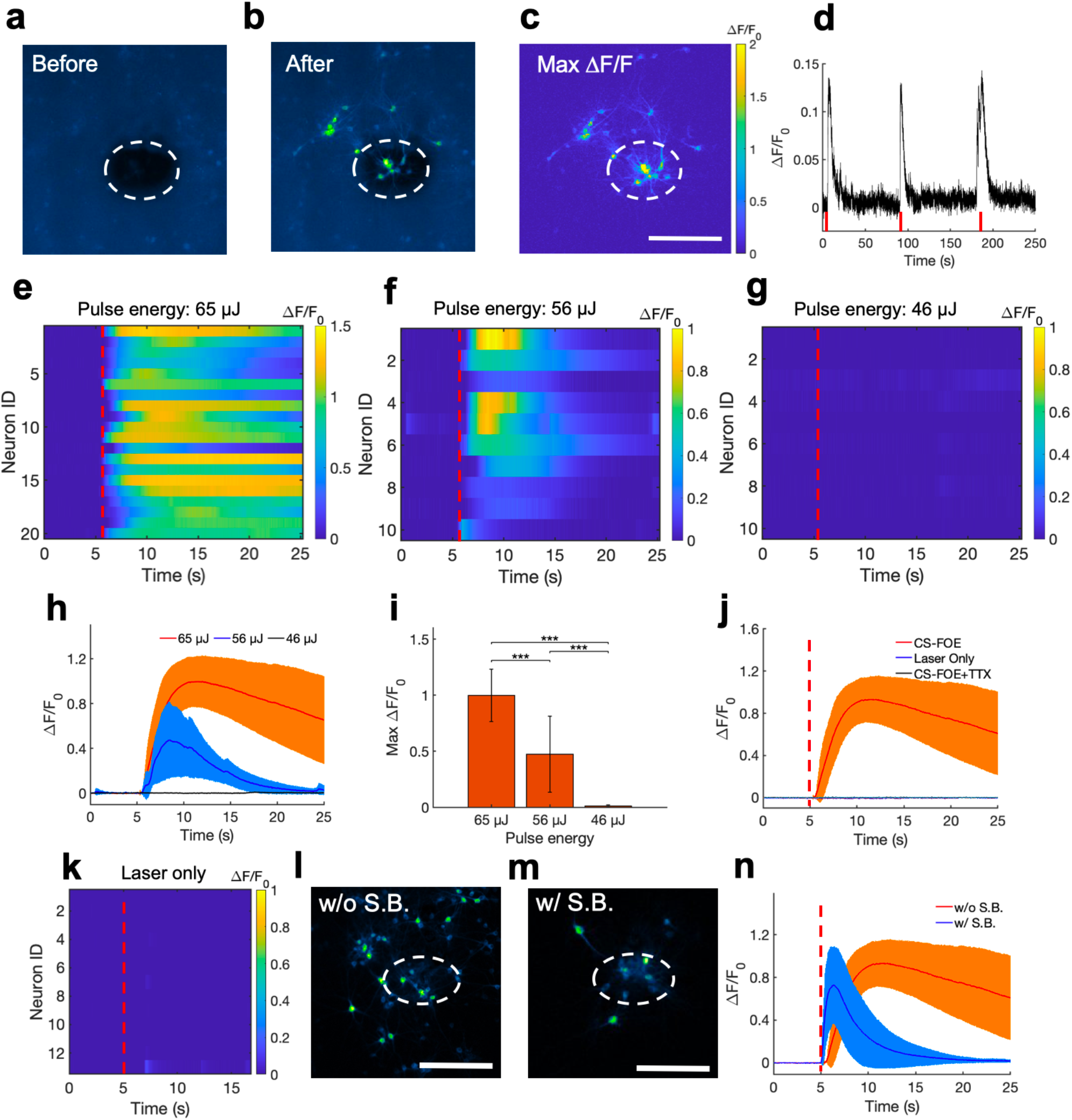
Activation of GCaMP6f-expressing cortical neurons by CSFOE stimulation. (a, b) representative fluorescence of neurons stimulated by CSFOE before stimulation (a) and after stimulation (b). (c) Map of the maximum fluorescence change ΔF/F_0_ induced by the CSFOE stimulation. Laser condition: 3 ms duration, pulse energy 65 μJ. Scale bar: 200 μm. (d) Calcium trace shows repeatable activation of the same neuron. Laser condition: 3 ms duration, pulse energy 56 μJ. (e-g) Colormaps of fluorescence change in neurons stimulated by CSFOE with a laser pulse energy of 65 μJ (e), 56 μJ (f), and 46 μJ (g). (h) Average calcium traces of neurons obtained from (e, f, g) with the pulse energy of 65 μJ (red), 56 μJ (blue) and 46 μJ (black), respectively. The shaded region corresponds to one standard deviation. Laser turns on at t = 5 s (red dashed lines). The duration of each stimulation was fixed at 3 ms. (i) Average of maximum fluorescence intensity changes shown in (e)-(g). Error bars represent standard deviation. (j) Average calcium traces of neurons of CSFOE stimulation, laser only control group, and TTX control group. (k) Colormaps of fluorescence change in neurons of a laser only control group. (l, m) Contrast calcium imaging of GCaMP6f transfected neurons without synaptic blocker (l) and with synaptic blocker (m). Scale bar: 200 μm. (n) Average calcium traces without (red) and with (blue) synaptic blocker. Laser turns on at t = 5s.

Next, we investigated the effect of laser pulse energy on CFOE stimulation. Each pulse train was fixed to be 3 ms long. Three laser pulse energies, 65, 56, and 46 μJ, were applied to the CSFOE to modulate neural activities. Responses from neurons at each pulse energy are plotted as heatmaps in **Figure 3e-g**. Representative calcium traces are plotted in **Figure 3h**. The averages of maximum fluorescence change obtained from these three groups are compared in **Figure 3i**. With the laser pulse energy of 65 and 56 μJ, neurons showed an average maximum fluorescence change (Δ*F*/*F*_0_) of 99.8 ± 23.3% and 47.4 ± 33.9%, while with laser energy of 46 μJ, the induced fluorescence change is negligible (1.2 ± 1.0%). These results indicate that at the laser pulse train with the repetition rate of 1.7 kHz and 3 ms duration, the activation threshold is between 46 μJ and 56 μJ, corresponding to a pressure of ∼8 MPa.

To confirm that the observed activation was due to optoacoustic stimulation, we performed a laser-only control and compared it to the calcium traces of CSFOE-stimulated neurons. The laser-only control group used the same optical fiber without any coatings on the tip with the same repetition rate of 1.7 kHz, 3 ms duration and laser pulse energy of 56 μJ. No significant fluorescence response was observed in the laser only group (**Figure 3j** and **k**). Optical excitation alone triggered negligible activities. Additionally, to investigate whether the activations observed were caused by action potential, we performed a control experiment with addition of 3 mM tetrodotoxin (TTX), a blocker of voltage-gated sodium channels. No significant fluorescence response was observed in the TTX group (**Figure 3j**), indicating that the observed calcium transients under CSFOE stimulation were induced by the firing of action potentials. These results are also consistent with previous studies of optoacoustic stimulation (Jiang et al., 2021).

To investigate how synaptic inputs affects the stimulation outcomes, we applied a cocktail of synaptic blockers (10 mM NBQX, 10 mM gabazine, and 50 mM DL-AP5). As shown in **Figure 3l** (and **Figure 3c**), when there was no synaptic blocker added, due to the network effect, many neurons outside the stimulation area were activated. With synaptic blocker added (**Figure 3m**), most of the stimulation effects were confined within the sub-millimeter area centered around the CSFOE. Averaged traces of stimulated neurons with and without synaptic blockers in both conditions are plotted in **Figure 3n**. Two types of neuron responses were observed: a transient response under synaptic blocking (blue) and a prolonged response without synaptic blocking (orange). The decay portion of the response curves can be fitted exponentially and a time constant for the decay can be defined at the time when the fluorescence intensity decreased by a factor of 1/e from the peak fluorescence intensity. The time constant decreased significantly from 12 s without synaptic blocking to 4 s with synaptic blocking. These results demonstrate that transient stimulation is likely the result of direct CSFOE optoacoustic stimulation, while the network effect through synaptic transmission results in prolonged stimulations (Jiang et al., 2021).

### 3.4 Comparison between CSFOE and FOC

To evaluate the performance improvement of CSFOE from the previous FOC fabricated using graphite and epoxy, we first compared the design of CSFOE and FOC. As shown in **Figure 4a**, both CSFOE and FOC have two-layer structures. Compared with FOC, several improvements were made on the CSFOE regarding the choice of material and structure design. Instead of using a graphite-epoxy system, a CS/PDMS mixture was used in CSFOE as the optoacoustic material. Compared to the previous design, CS has stronger absorption while PDMS is well known for its huge expansion coefficient of 310 μm m^-1^ C^-1^. The thickness of the CS layer in the CSFOE was optimized to obtain the largest pressure.

**Figure 4.**
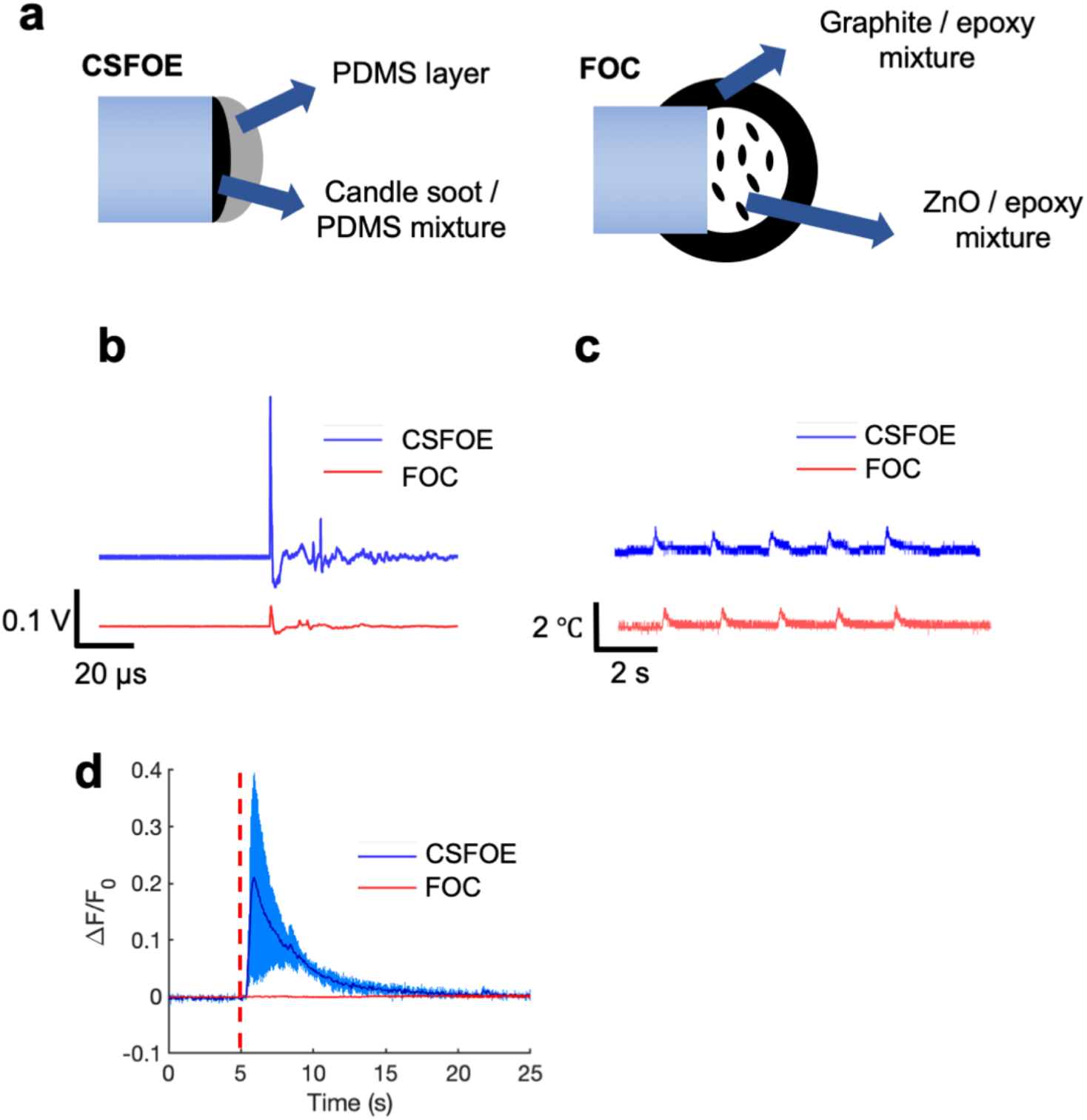
Comparison of CSFOE and FOC. (a) Schematics of the CSFOE and FOC. (b) Photoacoustic signal of CSFOE and FOC, measured by a 5 MHz transducer under the same laser condition: 1030 nm, 3 ns, 1.7 kHz, 48 mW. (c) Temperature rise measured by a thermal probe placed at the surface of CSFOE and FOC, respectively. (d) Representative calcium traces of GCaMP6f transfected neurons stimulated by CSFOE (Blue) and FOC (Red) under the same laser energy input of 52 μJ.

To directly compare the performance, we compared the pressure generated by CSFOE and FOC under the same laser condition. A transducer with greater sensitivity compared with the hydrophone was used to measure the generated pressure. As shown in **Figure 4b**, under the same laser condition of 1030 nm, 3 ns, 1.7 kHz, 48 mW, CSFOE generated a 9.6 times higher signal than that generated by FOC. In addition, the temperature rise associated with the optoacoustic conversion was measured for both fiber emitters using a thermal coupler placed on the surface the fiber tips. According to **Figure 4c**, the average temperature increases were 0.79 °C for the CSFOE and 0.77 °C for the FOC. Similar temperature increases suggest that while the CSFOE significantly increased the output pressure, the thermal effect remained minimal. Notably, both temperature increase was less than 1 °C, which is far below the threshold for photothermal neuron stimulation(Zhu et al., 2022). Such a small temperature increase also minimizes the risk of thermal damage for the neural system.

To compare their performance in neuron modulation, CSFOE and FOC were tested in GCaMP labelled neuron culture. Under the the laser condition of 3 ms pulse train, 56 μJ pulse energy, 1030 nm, 1.7 kHz repetition rate, successful activation was observed when CSFOE was applied to neurons. The average maximum Δ*F*/*F*_0_ reached over 20%. When FOC was applied under the same laser condition, no obvious activation occurred (**Figure 4d**). Notably, previous work showed Ca imaging signals indicating successful activation by FOC has been confirmed in Oregon Green labelled neuron culture. GCaMP and Oregon Green, as calcium sensors, have different sensitivity upon stimulation. It has been reported that for a single action potential, Oregon Green can generate ∼50% fluorescence change, while GCaMP6f can only generate ∼10%(Palmer et al., 2014;Dana et al., 2019). Collectively, our result clearly shows that CSFOE has a significantly higher stimulation efficacy and can be more widely used for recording based on different kinds of calcium sensors.

### 3.5 Dual-site neuron stimulation by CSFOE

To illustrate the advantage of the high optoacoustic conversion efficiency of CSFOE, we used CSFOE for dual-site neuron stimulation in vitro. A 1 × 2 fiber splitter was used for splitting the laser energy into two identical paths. The laser pulse energy of each path was 56 μJ (**Figure 5a**). As shown in **Figure 5b**, the map of maximum fluorescence changes Max ΔF/F_0_ clearly shows two groups of neurons, with each centered around a CSFOE, being successfully activated by two CSFOEs with fluorescence increase of around 10% at each site. Each group is confined within an area of ∼200 μm associated with the corresponding CSFOE. The highly localized feature of CSFOE stimulation makes it possible to distinguish different sites of stimulation under the same field of view.

**Figure 5.**
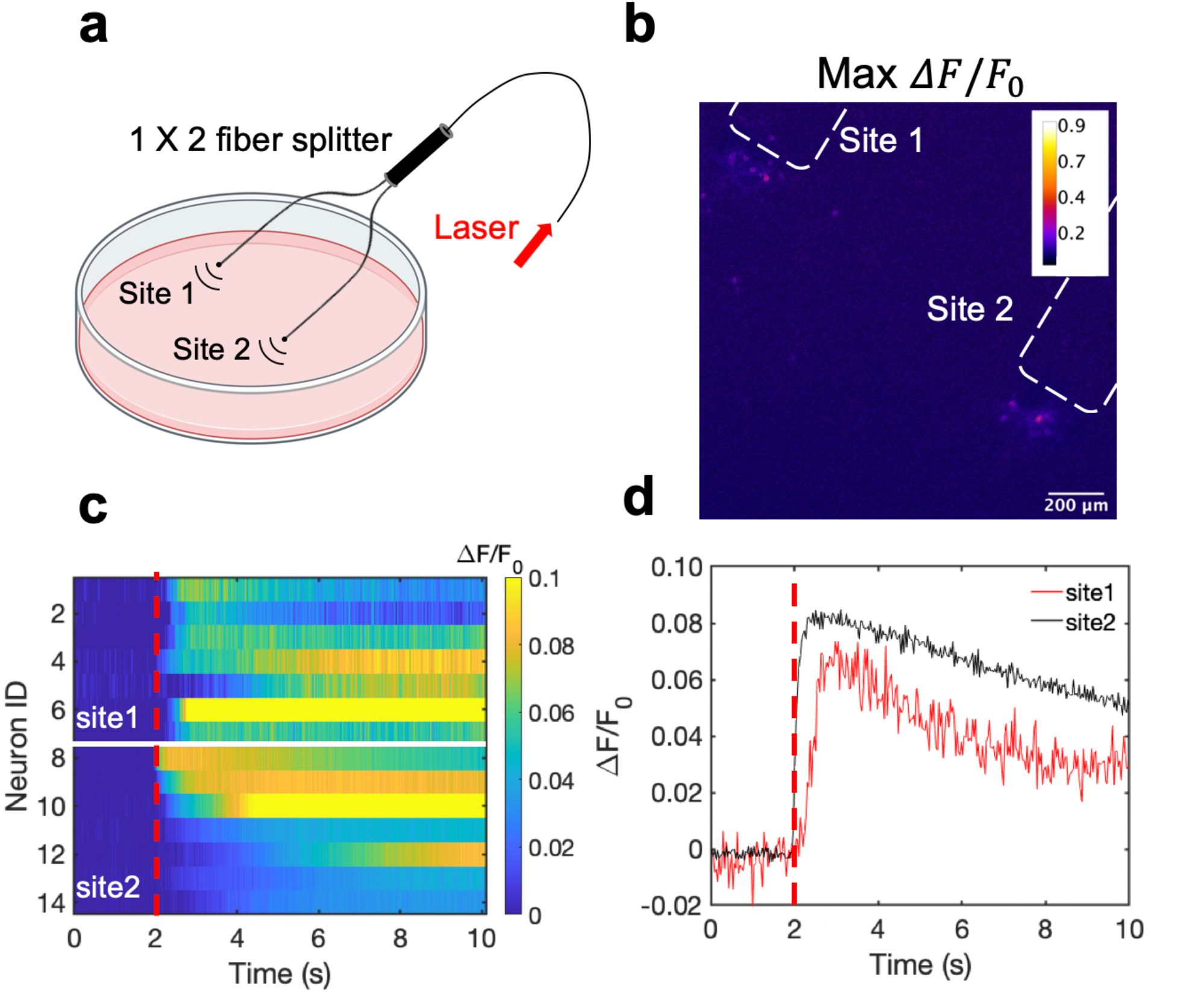
Dual site neuron stimulation by CSFOE. (a) Schematic of dual site stimulation using two CSFOEs with a fiber splitter. (b) Map of the max Δ*F*/*F* image of two sites of neurons stimulated by two CSFOE. (c) Colormaps of fluorescence changes in neurons at two sites stimulated by CSFOE. (d) Representative calcium traces of neurons at site 1 (red) and site 2 (black).

Ca traces from two groups at these two sites are plotted in a heatmap shown in **Figure 5c**. Representative traces of different sites are plotted in **Figure 5d**. Neurons in both sites showed significant change in fluorescence after the laser onset at t = 2 s. The fluorescence changes at each site all reached over 10%, which shows that both sites are successfully stimulated (**Figure 5c**). The high optoacoustic conversion efficiency and the highly localized stimulation area open up potentials for multi-site neuron stimulation.

## 4 Conclusion

In this study, we developed a new fiber optoacoustic emitter based on CS for the first time with high optoacoustic conversion efficiency and demonstrated CSFOE neuromodulation with an improved efficacy compared to FOC. Based on these improvements, we demonstrated dual-site neuromodulation through two CFOE driven by a single laser source.

To obtain the highest optoacoustic pressure, we chose candle soot as the material of the absorber, which is considered as one of the best materials for optoacoustic generation owing to its high optical absorption. In addition, we optimized the layered design of the CSFOE through both simulation and experiment. The optimized CSFOE was able to generate over 15 MPa peak-to-peak pressure. A more detailed comparison of photoacoustic conversion efficiency between CSFOE and other two fiber optoacoustic emitters used in neuromodulation is shown in Table 1 below.

**Table 1.** optoacoustic conversion efficiency comparison of different fiber based optoacoustic emitters for neuromodulation.

|  | CSFOE (This work) | TFOE (Shi et al., 2021) | FOC (Jiang et al., 2020) |
| --- | --- | --- | --- |
| Energy conversion efficiency (%) | 1.5E-3 | 2.28E-6 | 3.14E-5 |
| Optoacoustic conversion efficiency in pressure ($\text{Pa m}^2/\text{J}$) | 15600 | 130 | 1374 |

Through the direct comparison, CSFOE is ∼ 100 times more efficient than TFOE. Besides, CSFOE shows ∼10 times higher conversion efficiency compared with FOC, which is also evident in the results shown in **Figure 4b**.

Detailed optoacoustic characterization for CSFOE has also been performed, including power-pressure dependence and distance-pressure dependence. The output optoacoustic peak to peak pressure is linearly proportional to the input pulse energy as P = 0.226 * E. The distance-pressure dependence confirmed a highly localized ultrasound field of around 300 μm. Based on the results of optoacoustic characterization, we can precisely control the ultrasound intensity to be delivered to neurons by controlling the energy of the laser as well as the distance between CSFOE and neurons.

Successful CSFOE neuron activation has been demonstrated using Calcium imaging. It was found that under the pulse energy of 56 μJ and 65 μJ, at the repetition rate of 1.7 kHz, over a 3 ms duration, the maximum fluorescence change of the stimulated neurons were 47.4 ± 33.9% to 99.8 ± 23.3%, respectively. These laser conditions correspond to optoacoustic pressure of 8.8 MPa and 12.4MPa at the peak of frequency of 12.8 MHz for CSFOE.

Taking advantage of its high energy conversion efficiency, we performed the dual-site neuron stimulation using two CSFOEs driven by a single laser, which is not feasible by previous fiber based optoacoustic emitters. Dual-site stimulation has lots of potential applications in animal behavior studies, since complex animal behavior is normally controlled by multiple functional area in the brain. CSFOE, offering a superior sub-millimeter spatial resolution and high-pressure conversion efficiency, has the potential to modulate more complex animal behavior by controlling multiple target sites in the circuitry.

In summary, this robust and highly efficient optoacoustic converter, with an easy and repeatable fabrication process, offers a new tool for effective neuron stimulation. With an improved efficiency and the ability to perform multi-site stimulation, CSFOE opens up a great potential for complex animal behaviors that needs multiple stimuli at different locations in a programmable manner.

### Data Availability Statement

The original contributions presented in this study are included in the article, further inquiries can be directed to the corresponding author/s.

### Ethics Statement

All experimental procedures have complied with all relevant guidelines and ethical regulations for animal testing and research established and approved by the Institutional Animal Care and Use Committee of Boston University (PROTO201800534).

### Author Contributions

GC, LS, JXC and CY: drafting and refining the manuscript. GC: conducting of the simulation. GC and LS: conducting of the experiments. LL, JXC and CY: critical guidance of the project. RW, YL, ZD, MH: help with experiments. LL, YL and MH: critical reading of the manuscript. All authors have read and approved the manuscript.

### Funding

This work is supported by Brain Initiative R01 NS109794 to J.-X.C. and C.Y. by National Institute of Health, United States. We also thank Y. Tian for help with the neuronal cultures.

### Publisher’s Note

All claims expressed in this article are solely those of the authors and do not necessarily represent those of their affiliated organizations, or those of the publisher, the editors and the reviewers. Any product that may be evaluated in this article, or claim that may be made by its manufacturer, is not guaranteed or endorsed by the publisher.

